# Characterization of the habitat- and season-independent increase in fungal biomass induced by the invasive giant goldenrod and its impact on the fungivorous nematode community

**DOI:** 10.1101/2020.09.16.274845

**Authors:** Paula Harkes, Lisa J.M. van Heumen, Sven J.J. van den Elsen, Paul J.W. Mooijman, Casper W. Quist, Mariëtte T. W. Vervoort, Gerrit Gort, Martijn H.M. Holterman, Joris J.M. van Steenbrugge, Johannes Helder

## Abstract

Outside its native range, the invasive plant species giant goldenrod (*Solidago gigantea*) has been shown to increase belowground fungal biomass. This non-obvious effect is poorly characterized; we don’t know whether it is plant developmental stage-dependent, which fractions of the fungal community are affected, and whether it is reflected in the next trophic level. To address the questions, fungal assemblages in soil samples collected from invaded and non-invaded plots in two soil types were compared. Whereas ergosterol as a marker for fungal biomass demonstrated a significant increase in fungal biomass, specific qPCR assays did not point at a quantitative shift. MiSeq-based characterization of the belowground effects of giant goldenrod revealed a local increase of mainly Cladosporiaceae and Glomeraceae. This asymmetric boost in the fungal community was reflected in a specific shift in the fungivorous nematode community. Our findings provide insight in the potential impact of invasive plants on local fungal communities.

## Introduction

Deliberately or by accident, humans have been transferring plants all over the world for centuries. Most of the time, introduced plants will not invade native ecosystems because they are sub optimally equipped for the new environment. Exotic plants are considered naturalised once they are able to sustain self-replacing populations for at least ten years in their new non-native growth area (Pyšek et al. 2004). A subset of naturalised plants is able to spread widely and may reach high densities in their new environment. Such plants are referred to as invasive plants (Pyšek et al. 2004; Richardson et al. 2000) and can have a major impact on the invaded ecosystem’s structure and processes (Vilà et al. 2011).

Aboveground observations have often shown that invasive plants induce a decrease in species richness of the native plant community (Hejda et al. 2009). Belowground, invasive plants can change physical conditions and the composition of soil biota. Japanese barberry (*Berberis tunbergii*) is an example of an invasive shrub that changed the local soil function due to its easily degradable litter, which has a high nitrogen content (Ehrenfeld et al. 2001). A second example is the Australian legume *Acacia dealbata* that forms densely patches in Northwestern Spain. In various ecosystems, the presence of this invasive plant resulted in a local increase in N, exchangeable P and overall organic matter content (Lorenzo et al. 2010). Exotic plant species have also been reported to induce changes in soil microbial community. *Chromolaena odorata*, a perennial herb from Mexico that became highly invasive in China, gave rise to a local increase in fungal biomass (Xiao et al. 2014). Comparable changes were observed for *Solidago gigantea* and *Solidago canadensis*, two *Solidago* species from Northern America that established throughout Europe and Asia. *S. gigantea* was shown to boost the local fungal community (Quist et al. 2014), whereas *S. canadensis* was demonstrated to induce a qualitative change in the local soil fungal community (Wang et al. 2018).

Invasive plants may even negatively affect the soil biological conditions for the native plant community, rendering the restoration of the original vegetation more difficult. The non-mycorrhizal *Brassica nigra* (black mustard), is invasive in North America and was shown to negatively affect mycorrhizal symbiosis. Thereby, making it more difficult for mycorrhizal plants - the vast majority - to establish in its vicinity (Pakpour and Klironomos 2015). Similarly, the non-mycorrhizal garlic mustard (*Alliaria petiolata*), an invasive species in North American forests, has a strong negative effect on native mycorrhizal communities, whereas in its native range (Europa) this effect is much milder (Callaway and Ridenour 2004). On the contrary, the mycorrhizal *S. canadensis* releases secondary metabolites in the rhizosphere that promote the growth of its own arbuscular mycorrhiza in the invaded area (Yuan et al. 2014).

To study belowground effects of invasive plants, it is advantageous to select rhizomatous perennial herbs. Rhizomes are subterranean stem parts that give rise to new stems. This mode a vegetative reproduction gives rise to dense, genetically uniform stands. Perennials are preferred as shifts in microbial communities might accumulate over years (Harkes et al. 2017). Giant goldenrod (*S. gigantea;* Asteraceae) is rhizomatous perennial herb native to North America (Weber and Jakobs 2005). After its introduction in Europe as an ornamental in the 18^th^ century (Weber 1998), it became a widespread invasive plant. *S. gigantea* can survive under a broad range of light intensities, soil moistures, temperatures, nutrient conditions and pH (Vanderhoeven et al. 2006). In its natural range, *S. gigantea* is colonized by mycorrhizal fungi (Wardle et al. 2004). (Zubek et al. 2016) showed that giant goldenrod interacts with AMF outside its native range, and the mycorrhizal frequency was higher in invaded as compared to neighboring non-invaded plots. Being a rhizomatous perennial herb that forms well-nigh monoculture stands in various habitats, giant goldenrod is an auspicious species to study the effect of invasive plants on soil biota.

In previous studies on the belowground effects of the invasive giant goldenrod, a local increase in the overall fungal biomass was detected, both in a mesocosm experiment (Scharfy et al. 2010), and under semi-natural conditions (Quist et al. 2014; Stefanowicz et al. 2016). The total fungal biomass was assessed by ergosterol, a biochemical marker for higher fungi, or by PLFA 18:2ω6. Ergosterol is a valid marker for major fungal groups such as Ascomycota and Basidiomycota, but is should be noted that some fungal groups such as the Glomeromycota and the Chytridiomycota lack this sterol in their cell membranes (see *e.g.* (Weete et al. 2010)).

Invasive plant-induced changes in the fungal community might be mirrored among fungivorous metazoan commity. Fungivorous nematodes are informative in this context as they are present at high densities in nearly any soil habitats, and as their ability to feed on fungi arose multiple times independently (Holterman et al. 2017) resulting in lineages with distinct preferences (Baynes et al. 2012; Okada and Kadota 2003). Previously, it was shown that a giant goldenrod-induced boost in fungal biomass was translated into an increase of a subset of the fungivorous nematode lineages (Quist et al. 2014).

The relevance of soil type and location for the impact of *S. gigantea* on fungal biomass was underlined by (Stefanowicz et al. 2016). They investigated 16 *S. gigantea-invaded* sites with two adjacent paired-plots (2 m x 2 m) at each site either in or outside a river valley. A local increase of fungal phospholipid fatty acids (PLFA) was observed in the *S. gigantea-invaded* plots, and this effect was more prominent in areas next to the river - directly exposed to fluvial processes - than in the areas just outside the river valley.

Here we investigated the impact of invasive *S. gigantea* on local fungal communities in more detail. First, we verified whether the *S. gigantea-induced* increase in fungal biomass was transient or long lasting. Therefore, soil samples were collected at the end of the growing season (November), whereas (Quist et al. 2014) mapped this phenomenon in September, and (Stefanowicz et al. 2016) in August. Second, ribosomal DNA-based markers were used next to ergosterol to characterize changes in the fungal community. The use of two independent markers for fungal biomass could provide a more solid basis for our findings, and fungal division-specific markers would allow us to characterize the impact qualitatively. Whereas the biomass marker ergosterol pointed at a stimulation of at least a major part of the fungal community, a general rDNA marker for fungi as well as markers for major constituents of the fungal community showed no effect of the presence of invasive *S. gigantea*. To further investigate these apparently contradictory results, ribosomal DNA amplicons were sequenced in order to investigate which fungal families were indicative for invaded plots. In addition, we checked whether representatives of the next trophic level, fungivorous nematodes, were affected. Two out of the three nematode lineages present on these sites were stimulated in the presence giant goldenrod. Possible explanations for these interesting but paradoxical results are discussed.

## Material and Methods

### Sampling sites

The belowground *S. gigantea* invasion effects were examined at eight sites in the Netherlands, located in either of the distinct semi-natural habitats, namely riparian zones (rive clay soil) and semi-natural grasslands (sandy soil). To allow for a comparison with results presented by Quist et al. (2014), the same sampling sites were used. ‘Millingerwaard’, ‘Ewijkse Plaat’ and ‘Blauwe Kamer’ were the selected riparian zone sites (Table 1). The other five sites, ‘Dennenkamp’, ‘Plantage Willem III’, ‘Hollandseweg’, ‘Scheidingslaan’ and ‘Reijerscamp’, are located in semi-natural grassland habitats on Pleistocene sandy soils (Table 1). For all invaded plots, the coverage by *S. gigantea* was scored as a 9 on a modified Braun-Blanquet scale (Barkman et al. 1964; Leps and Hadincova 1992) implying a 75-100% coverage. Non-invaded plots were dominated by native plant species and fell in category 2 which means that at most 2-5 *S. gigantea* were found in the control plots. More information on the floristic composition of these sites can be found in (Quist et al. 2014).

**Table 1:**
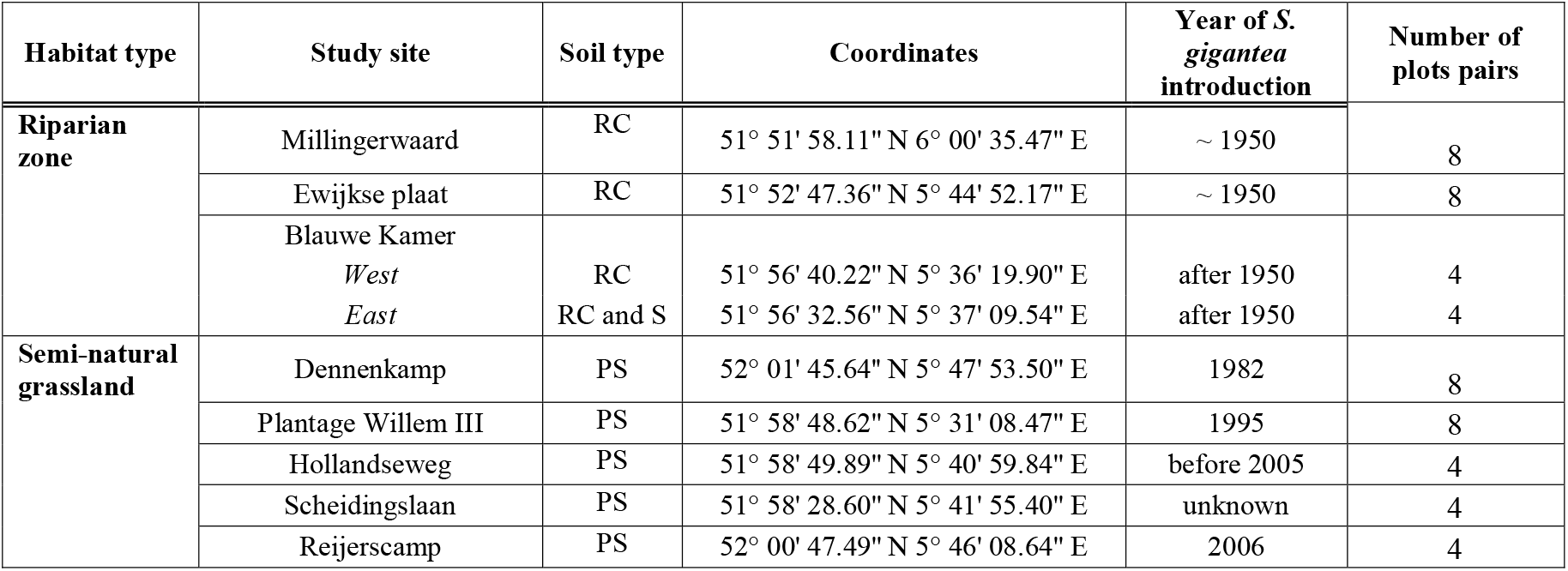
Eight study sites located in riparian zones and in semi-natural grassland habitats are indicated below. RC = River clay, S = Sand and PS = Pleistocene sand. Although ‘Blauwe Kamer’ is one riparian study site, samples were collected from two distinct areas within the nature reserve (1 and 2). Riparian zones are characterized by river clay soils, whereas the semi-natural grassland sites were located on Pleistocene sandy soils. Coordinates and years of *S. gigantea* introduction were obtained from Quist et al. (2014)

### Soil sampling

In total, 104 composite soil samples were collected from 52 plot-pairs in November 2014. Eight plot-pairs were selected per site for Millingerwaard, Ewijkse Plaat, Blauwe Kamer, Dennenkamp and Plantage Willem III. Four plot-pairs were sampled at the sites Hollandseweg, Scheidingslaan and Reijerscamp due to limited number of *S. gigantea* patches at these sites (see also Table 1). Each plot-pair consisted of two directly neighbouring 2 x 2 m plots to minimize possible differences in soil type and structure. To average microscale variation, 12 soil cores (depth: 25 cm, ø 1.5 cm) were randomly collected within each plot and mixed thoroughly. Sampling material was thoroughly cleaned between plot-pairs in order to limit cross contamination. At the day of sample collection composite soil samples were split into two subsamples (200g and 5g). The 200g subsample was stored at 4°C for subsequent nematode extraction (100g) and the determination of abiotic soil characteristics (60g). Nematodes were extracted within one week after sample collection. The other subsamples (5g) were stored at −20°C to prevent DNA degradation prior to total DNA extraction, which was completed within three weeks after sample collection.

### Abiotic soil characteristics

Per composite sample, subsamples were taken for the analysis of abiotic and biotic soil characteristics. Moisture content, pH, organic matter (OM) content, total carbon (C) content, total nitrogen (N) content and C:N ratio were determined. The total amount of C and N, determined with a composite sample of invaded and a composite sample of uninvaded plots per sampling site, was performed by BLGG AgroXpertus (Wageningen, The Netherlands).

Soil moisture content was measured per sample by determining the weight loss after 20 hours at 105°C. Dried soil was sieved with a mesh of 2 mm and 10 g was added to 25 ml demineralized water for soil pH measurements using a gel-electrolyte electrode (Sentix 21, WTW, Weilheim, Germany). Organic matter content was determined by measuring weight loss of 20 g of sieved soil after 5 hours at 550°C.

### Fungal and bacterial extraction and community analysis

Fungal and bacterial DNA was extracted from 0.25 g subsamples using the PowerSoil DNA Isolation Kit (MO BIO Laboratories, Carlsbad, California, USA). Slight changes were made to the manufacturer’s protocol. PowerBead Tubes were placed in a Qiagen Tissue Lyser for 7 minutes instead of 10 minutes to compensate for the high shaking frequency (30 Hz), and 20 μl of internal control DNA (20.6 ng/μl calf thymus DNA) was added to each sample in order to monitor DNA losses during extraction and purification. To further reduce the impact of soil-derived PCR-inhibiting components, purified lysates were diluted 100 times. Diluted samples were stored at 4 °C until further use. Undiluted purified samples were stored at −20 °C. Microbial communities were analysed using real time PCR assays targeting total fungi (ITS1F/5.8 s), total bacteria (16S ribosomal RNA) as well as three fungal phyla: Ascomycota, Basidiomycota and Chytridiomycota (based on taxon-characteristic ITS regions). PCR primers, PCR conditions, and slope and intercept values describing the relationship between Ct-values and concentration of target bacterial or fungal DNA (ng/μl) are essentially according to (Harkes et al. 2017) and details can be found in Supplementary table 1.

### Ergosterol measurements

Ergosterol, a biochemical marker for higher fungi frequently used in soil ecology, was extracted from 1 g of soil using the alkaline extraction method as described by (Bååth 2001). In a mixture alkaline methanol and cyclohexane, ergosterol accumulated in the cyclohexane phase. After phase separation, the cyclohexane was removed by evaporation, and ergosterol is re-dissolved in methanol. Subsequently, high-performance liquid chromatography (HPLC) with photodiode array detection (peak identification is based on retention time and UV-spectrum) was used to separate and quantify the ergosterol contents of the samples as described by (de Ridder-Duine et al. 2006)

### Nematode extraction and community analysis

Per composite sample, nematodes were extracted from a 100g subsample using an Oostenbrink elutriator (Oostenbrink 1960). DNA extractions of the total nematode suspensions were performed as described by (Vervoort et al. 2012). At the start of this extraction procedure, 25 μl of calf thymus DNA (20.6 ng/μl) was added to each sample to be able to quantify DNA loss after extraction and purification. After purification, each sample was diluted 10 times and stored at −20°C until further use. Diluted DNA extracts served as a template for the real time PCR-based determination of the total nematode density and the densities of the three major fungivorous nematode lineages present on these locations, Aphelenchidae, Aphelenchoididae and *Diphtherophora*. This qPCR detection method is based on taxon-specific SSU rDNA sequence motifs as previously described by (Vervoort et al. 2012).

Because of the substantial variation estimation in rDNA copy numbers in fungi, using only the ITS marker might not suffice in all fungal clades, therefore two qPCR assays for single-copy protein coding genes were included in this study (beta-tubulin (tub2) and translation elongation factor 1-alpha (tef1)) as they are supposed to be less viarable and occure as a single copy in fungi (Raja et al. 2017).

### PCR Amplification and sequencing of fungal 16S rDNA

The variable V7-V8 of fungal 18S was utilized as a target for the analyses of Illumina 18S rDNA sequencing. To prepare the samples for sequencing a twostep PCR procedure was followed as described in (Harkes et al. 2019). In brief, a locus-specific primer combination extended with an Illumina read area and the appropriate adapter were used to produce primary amplicons - in triplicate for all samples. PCR 2 was conducted on 40x diluted amplicons of PCR1 to attach the Illumina index and the Illumina sequencing adaptor. Randomly picked products of PCR 1 and 2 were checked on gel to ensure amplification was successful. Finally, all PCR products were pooled and sent for sequencing. Sequencing was done at Bioscience - Wageningen Research, Wageningen, The Netherlands - using the Illumina MiSeq Desktop Sequencer (2*250nt paired-end sequencing) according to the standard protocols. The raw sequences were submitted to the NCBI Sequence Read Archive (SRA) database under study accession numbers PRJNA563313.

### Combined analysis of abiotic characteristics and quantitative biotic data

The impact of *S. gigantea* invasion on abiotic soil properties and the densities of nematodes, fungi and bacteria was analysed by using mixed linear models (PROC MIXED, SAS software system version 9.2, see Littell et al. (2006). When residuals did not approximate normal distributions, transformed data were used. OM, total C, total N, nematode densities and densities of fungi and bacteria were log-transformed. A constant of 0.1 was added prior to the log-transformation to bypass any zero values. This was done for Aphelenchidae, Aphelenchoididae, *Diphtherophora* and Chytridiomycota.

A split-plot design was used for all ten study sites, with sampling sites forming the main plots, associated with the factor habitat type (riparian vegetation or semi-natural grassland), with multiple plot pairs (8 or 4) per site, and two subplots per plot pair, associated with the factor plant invasion. This design was represented in the mixed models with random effects for sites, plot-pairs and individual plots, forming the random part of the model. Main effects of habitat type, invasion and the interaction between both factors formed the fixed part of the model. Random effects for site, plot-pairs and individual plots formed the random part of the model. In this way, the total error variance was split into variance components for sites, plot-pairs within sites and for individual plots within plot-pairs. Regarding pH, the mixed model took into account that variances were different for riparian vegetation habitats and semi-natural grasslands (as was noticed from residual plots). Hypothesis tests (with F-test statistics) for the significance of the main effects of habitat type, invasion and their interaction on the soil variables were performed. P-values <0.05 were considered significant. Regardless the outcome of hypothesis tests on interaction and main effects, comparisons between invaded and uninvaded plots were made per habitat type, using F-tests. The results were presented as (back transformed) 95% confidence intervals for the estimated mean responses of the soil variables (obtained from ‘least squares means’ outputs) in invaded and un-invaded plots per habitat type. Moreover, invasion impacts on soil variables were presented as ratios between estimated means of invaded plots and un-invaded plots.

### Bioinformatics framework and statistics

The composition of the fungal communities of the soil samples was analysed based on the sequencing data obtained from the Illumina MiSeq platform. Reads were sorted into the experimental samples according to their index combination, quality trimmed by BBDUK and then merged via VSEARCH (Bushnell 2018; Rognes et al. 2016). Unique sequences were then clustered at 97% similarity by using the usearch_global method implemented in VSEARCH and a representative consensus sequence per *de novo* OTU was determined (Rognes et al. 2016). The clustering algorithm also performs chimera filtering to discard likely chimeric OTUs with UCHIME algorithm in *de novo* mode (Edgar et al. 2011) implemented in VSEARCH. Sequences that passed quality filtering were then mapped to a set of representative consensus sequences to generate an OTU abundance table. Representative OTU sequences were assigned to a taxonomic classification via BLAST against the Silva database (version 12.8). Sequences not belonging to fungi were discarded from the 18S fungal dataset. Low-abundance OTUs (those with an abundance of <0.005% in the total data set) were discarded (Bokulich et al. 2013) prior to analysis. Samples were transformed using Hellinger transformation for all downstream analyses.

To investigate the indicator taxa involved in the differences in fungal communities between invasive and un-invasive, a linear discriminate analysis (Gonzalez et al.) effect size (LEfSe) was conducted in Microbiome Analyst (Dhariwal et al. 2017) to explore the differential microbial populations at family level (Segata et al. 2011). A significance level of α□≤□0.05 was used in this study.

## Results

### Changes in abiotic soil characteristics upon *S. gigantea* invasion

To gain insight in the abiotic environment of the *S. gigantea-invaded* sites, the soil moisture content, pH, OM content, total C content, total N content and C:N ratio were analysed. Significant changes were observed in soil moisture content, pH and OM content between *S. gigantea-invaded* and un-invaded plots in riparian and semi-natural grasslands sites (Table 2). In contrast, no differences were observed between invaded and un-invaded plots for the total C content, total N content and the C:N ratio (Tables 2 and 3, Figure 1).

**Figure 1.**
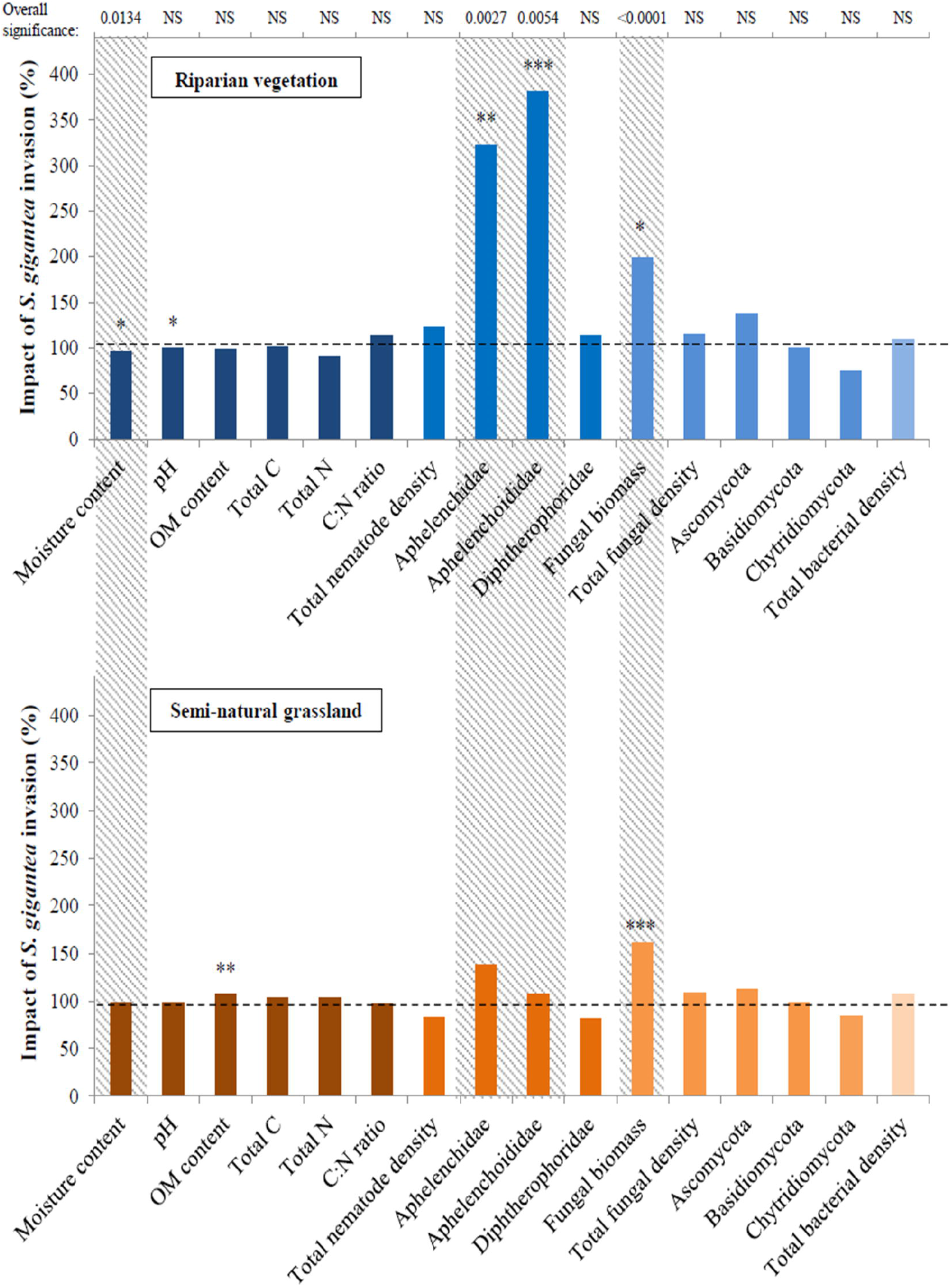
Impact of *S. gigantea* invasion in riparian vegetation (top) and semi-natural grassland habitat. The impact of *S. gigantea* invasion on the y-axis was calculated by dividing estimated means (Table 3) from invaded plots by estimated means from un-invaded plots and expressed as a percentage. Impacts are shown for the 6 abiotic variables, the total nematode density, densities of three fungivorous nematodes, total fungal density, densities of three fungal phyla and the total bacterial density (no invasion impact = 100%). Asterisks indicate significant differences (*P<0.05, **P<0.01, ***P<0.001) between invaded and un-invaded plots per habitat type. Variables showing an overall significant invasion effect, for both habitats together, are indicated by a grey-shaded background. Corresponding P-values are shown at the top part of the figure (NS=not significant). Riparian vegetation habitats included 3 study sites and 24 plotpairs, while semi-natural grasslands included 5 study sites and 28 plot-pairs.

**Table 1.** Main effects of habitat type, invasion and their interaction for the different abiotic and biotic variables analysed. F-test F_df_ values and corresponding P-values obtained from the mixed models are shown for each variable. Total C and N contents are expressed in g/kg dry soil. Total nematode density, Aphelenchidae, Aphelenchoididae and *Diphtherophora* are expressed in numbers (#) per 100 g dry soil. Total fungal density, Ascomycota, Basidiomycota, Chytridiomycota and total bacterial density are expressed in μg DNA per 100 g dry soil. Fungal biomass expressed as mg ergosterol kg^-1^ soil. The degrees of freedom (Crowther et al.) for *Diphtherophora* are lower than for the other variables, since this taxon was not present at two study sites (Scheidingslaan and Reijerscamp). Regarding invasion and interaction effects, the df for total C, N and C:N ratio are lower since samples were pooled together per study site. Significant P-values (<0.05) are indicated in bold.

|  | Habitat type |  |  | Invasion |  |  | Habitat type * Invasion |  |  |
| --- | --- | --- | --- | --- | --- | --- | --- | --- | --- |
| Abiotic variables | $F_{df}$ | | P-value | $F_{df}$ | | P-value | $F_{df}$ | | P-value |
| Moisture content (%) | $F_{1,6}$ | 2.02 | 0.2052 | $F_{1,50}$ | 6.58 | **0.0134** | $F_{1,50}$ | 0.93 | 0.3391 |
| pH | $F_{1,6}$ | 92.22 | **<.0001** | $F_{1,50}$ | 2.48 | 0.1213 | $F_{1,50}$ | 0.19 | 0.6661 |
| OM content (%) | $F_{1,6}$ | 0.13 | 0.7343 | $F_{1,50}$ | 2.87 | 0.0965 | $F_{1,50}$ | 4.74 | **0.0341** |
| Total C (g/kg) | $F_{1,6}$ | 1.38 | 0.2848 | $F_{1,6}$ | 0.05 | 0.8281 | $F_{1,6}$ | 0.00 | 0.9562 |
| Total N (g/kg) | $F_{1,6}$ | 0.57 | 0.4803 | $F_{1,6}$ | 0.15 | 0.7090 | $F_{1,6}$ | 0.60 | 0.4692 |
| C:N ratio | $F_{1,6}$ | 0.73 | 0.4267 | $F_{1,6}$ | 0.43 | 0.5378 | $F_{1,6}$ | 1.00 | 0.3568 |
| **Biotic variables** |  |  |  |  |  |  |  |  |  |
| Total nematode density (#) | $F_{1,6}$ | 26.69 | **0.0021** | $F_{1,50}$ | 0.00 | 0.9758 | $F_{1,50}$ | 3.24 | 0.0780 |
| Aphelenchidae (#) | $F_{1,6}$ | 0.02 | 0.8962 | $F_{1,50}$ | 9.96 | **0.0027** | $F_{1,50}$ | 3.16 | 0.0814 |
| Aphelenchoididae (#) | $F_{1,6}$ | 0.65 | 0.4507 | $F_{1,50}$ | 8.44 | **0.0054** | $F_{1,50}$ | 6.92 | **0.0113** |
| *Diphtherophora* (#) | $F_{1,4}$ | 0.08 | 0.7930 | $F_{1,42}$ | 0.02 | 0.9007 | $F_{1,42}$ | 0.29 | 0.5945 |
| Fungal biomass (mg erg / kg) | $F_{1,6}$ | 3.63 | 0.1055 | $F_{1,48}$ | 21.97 | **<0.0001** | $F_{1,48}$ | 0.72 | 0.3990 |
| Fungal DNA ($\mu\text{g}$) | $F_{1,6}$ | 0.75 | 0.4184 | $F_{1,50}$ | 2.50 | 0.1203 | $F_{1,50}$ | 0.17 | 0.6846 |
| - Ascomycota ($\mu\text{g}$) | $F_{1,6}$ | 0.06 | 0.8187 | $F_{1,50}$ | 3.34 | 0.0738 | $F_{1,50}$ | 0.71 | 0.4027 |
| - Basidiomycota ($\mu\text{g}$) | $F_{1,6}$ | 10.83 | **0.0166** | $F_{1,50}$ | 0.00 | 0.9637 | $F_{1,50}$ | 0.01 | 0.9370 |
| - Chytridiomycota ($\mu\text{g}$) | $F_{1,6}$ | 1.07 | 0.3405 | $F_{1,50}$ | 1.25 | 0.2693 | $F_{1,50}$ | 0.12 | 0.7340 |
| Total bacterial density ($\mu\text{g}$) | $F_{1,6}$ | 0.05 | 0.8360 | $F_{1,50}$ | 3.29 | 0.0759 | $F_{1,50}$ | 0.02 | 0.8826 |

**Table 3.** Estimated mean response and associated 95% confidence intervals of the soil characteristics analyzed for plots invaded and un-invaded by *S. gigantea* in two habitat types. The estimated mean response (Est. mean) and lower and upper bounds of the 95% confidence interval are shown for each variable. Values were obtained from ‘least squares means’ outputs of mixed models fitted to the variables. For both habitat types, riparian vegetation and semi-natural grassland, est. mean responses are shown for plots invaded and un-invaded by *S. gigantea*. Riparian vegetation habitats contained 24 plot-pairs in total, while semi-natural grasslands contained 28 plot-pairs in total. Values for OM content, total C (g/kg dry soil), total N (g/kg dry soil), Aphelenchidae, Aphelenchoididae, *Diphtherophora*, total fungi, Ascomycota, Basidiomycota, Chytridiomycota and total bacteria were back transformed from logarithmic values to the original scale. Aphelenchidae, Aphelenchoididae and *Diphtherophora* are expressed in numbers (#) per 100 g dry soil. For *Diphtherophora*, 8 plot-pairs from semi-natural grasslands were excluded from analysis. Fungal biomass expressed as mg ergosterol per kg soil. Total fungal density, Ascomycota, Basidiomycota, Chytridiomycota and total bacterial density are expressed in μg DNA per 100 g dry soil. Significant P-values (<0.05) are indicated in bold.

|  | Riparian vegetation (n=24 plot-pairs) |  |  |  |  |  |  | Semi-natural grassland (n=28 plot-pairs) |  |  |  |  |  |  |
| --- | --- | --- | --- | --- | --- | --- | --- | --- | --- | --- | --- | --- | --- | --- |
|  | Invaded (n=24) |  |  | Un-invaded (n=24) |  |  | P-value | Invaded (n=28) |  |  | Un-invaded (n=28) |  |  | P-value |
|  | Lower | Est. mean | Upper | Lower | Est. mean | Upper |  | Lower | Est. mean | Upper | Lower | Est. mean | Upper |  |
| Abiotic variables |  |  |  |  |  |  |  |  |  |  |  |  |  |  |
| Moisture content (%) | 15.9 | 20.7 | 25.6 | 16.7 | 21.5 | 26.4 | 0.0199 | 12.8 | 16.6 | 20.4 | 13.2 | 17.0 | 20.7 | 0.2445 |
| pH | 7.33 | 7.46 | 7.59 | 7.37 | 7.50 | 7.63 | 0.0197 | 5.21 | 5.58 | 5.96 | 5.28 | 5.65 | 6.02 | 0.3059 |
| OM content (%) | 4.2 | 5.9 | 8.3 | 4.2 | 5.9 | 8.4 | 0.7429 | 4.3 | 5.7 | 7.4 | 4.0 | 5.3 | 6.9 | 0.0063 |
| Total C (g/kg) | 17.0 | 31.2 | 57.0 | 16.8 | 30.6 | 56.0 | 0.9181 | 13.9 | 22.1 | 35.3 | 13.5 | 21.5 | 34.4 | 0.8243 |
| Total N (g/kg) | 0.9 | 1.7 | 3.1 | 1.0 | 1.9 | 3.4 | 0.4895 | 0.9 | 1.5 | 2.3 | 0.9 | 1.4 | 2.2 | 0.7663 |
| C:N ratio | 12.9 | 19.1 | 25.2 | 10.6 | 16.8 | 22.9 | 0.3366 | 10.5 | 15.2 | 20.0 | 10.9 | 15.7 | 20.5 | 0.7879 |
| Biotic variables |  |  |  |  |  |  |  |  |  |  |  |  |  |  |
| Aphelenchidae (#) | 0.4 | 1.6 | 6.0 | 0.1 | 0.5 | 2.0 | 0.0015 | 0.4 | 1.2 | 3.6 | 0.3 | 0.9 | 2.7 | 0.3156 |
| Aphelenchoididae (#) | 1.4 | 4.3 | 13.0 | 0.3 | 1.1 | 3.6 | 0.0004 | 1.6 | 4.0 | 9.8 | 1.5 | 3.7 | 9.2 | 0.8404 |
| *Diphtherophora* (#) | 0.4 | 0.9 | 1.9 | 0.4 | 0.8 | 1.7 | 0.7621 | 0.4 | 0.9 | 2.0 | 0.5 | 1.1 | 2.4 | 0.6564 |
| Fungal biomass (mg erg / kg) | 1.07 | 1.75 | 2.85 | 0.54 | 0.88 | 1.43 | 0.0199 | 1.74 | 2.66 | 3.97 | 1.22 | 1.65 | 2.76 | <.0001 |
| Fungal DNA (µg) | 220 | 299 | 405 | 191 | 259 | 351 | 0.1814 | 264 | 340 | 438 | 243 | 313 | 403 | 0.3925 |
| - Ascomycota (µg) | 26.1 | 40.2 | 61.9 | 19.1 | 29.3 | 45.2 | 0.0748 | 26.9 | 38.7 | 55.6 | 24.0 | 34.5 | 49.5 | 0.4730 |
| - Basidiomycota (µg) | 1.6 | 2.8 | 5.1 | 1.6 | 2.8 | 5.0 | 0.9818 | 5.4 | 8.8 | 14.3 | 5.5 | 9.0 | 14.5 | 0.9270 |
| - Chytridiomycota (µg) | 0.4 | 0.9 | 1.8 | 0.6 | 1.2 | 2.4 | 0.3250 | 0.3 | 0.6 | 1.2 | 0.4 | 0.7 | 1.4 | 0.5709 |
| Total bacterial density (µg) | 3598 | 4929 | 6754 | 3303 | 4526 | 6201 | 0.1875 | 3654 | 4696 | 6034 | 3399 | 4367 | 5612 | 0.2263 |

Plots invaded by *S. gigantea* had a lower soil moisture content than un-invaded plots (F_1,50_= 6.58, *P*= 0.0134; Table 2, Figure 1). This overall effect could mainly be attributed to the slightly lower moisture content of invaded plots in the riparian vegetation habitats (F_1,50_= 5.79, *P*= 0.0199; Table 3, Figure 1).

Riparian vegetation habitats and semi-natural grasslands differed significantly in pH (F_1,6_= 92.22, P<0.0001; Table 2). Riparian vegetation sites had a slightly alkaline soil with pH 7.5, while semi-natural grasslands had a moderately acidic soil with pH 5.6 (see Table 3 for 95% confidence intervals). Overall, no effect of invasion of soil pH was detected (Table 2, Figure 1). Splitting by habitat type however, showed that for both types the pH was slightly lower in invaded plots, but only for riparian sites this difference was significant, due to the lower variance in riparian plots (F1,50= 5.81, P= 0.0197; Table 3).

A significant interaction between invasion and habitat was found for OM content (F_1,50_= 4.74, *P*=0.0341), indicating that the effect of invasion was habitat type-dependent (Table 2). In semi-natural grasslands, *S. gigantea*-invaded plots had a higher OM content as compared to un-invaded plots (F_1,50_= 8.12, *P*= 0.0063; Table 3, Figure 1), whereas no difference in OM content was detected between plot-pairs at riparian sites.

### Invasive *S. gigantea* increase fungal biomass, but not the total fungal DNA

Using ergosterol as a biochemical marker for biomass of higher fungi, a strong overall effect of *S. gigantea* was detected (F_1,48_= 21.97, *P* <0.0001; Table 2). In giant goldenrod-invaded plots, a significant increase in ergosterol levels was observed for both habitat types (Tables 2 and 3). It is noted that ergosterol is an important constituent of the cell membranes of higher fungi, and as such it correlates fairly well with fungal biomass (e.g. (Newell and Fallon 1991). Using real time PCR assays, the total bacterial and total fungal communities were assessed, and no significant differences in fungal and bacterial DNA concentrations were observed, neither between invaded and non-invaded habitats, nor between the two habitat types (Table 3). Also, the two single-copy fungal protein coding genes included in this study (tub2 and tef1) did not show any significant differences between invaded and uninvaded plots (data not shown).

Keeping in mind that ergosterol measurements predominantly reflect the presence of Ascomycota and Basidiomycota, representatives of two major distal clades within the kingdom Fungi (Weete et al. 2010), these phyla were quantified separately. In the riparian vegetation habitats, a trend was observed of Ascomycota having a higher DNA concentration in *S. gigantea-invaded* plots (F_1,50_=3.31, *P*=0.0748; Table 3, Figure 1). A similar invasion effect was observed when both habitats were analysed together (F_1,50_=3.34, *P*=0.0738; Table 2). It is noted that the mean DNA concentration of Basidiomycota on sandy soils was about three times higher than the DNA concentration in the river clay soils (F_1,6_=10.83, *P*=0.0166; Table 2). The DNA concentrations of Basidiomycota did not differ between invaded and un-invaded plots (Tables 2 and 3, Figure 1). In addition, Chytridiomycota were measured, being a fungal phylum that is uses cholesterol instead of ergosterol as its major sterol, but no differences were observed between giant goldenrod-invaded and non-invaded plots. Comparison of Chytridiomycota between the two major habitats revealed no difference in DNA concentrations.

The overall bacterial DNA concentration tended to be slightly higher in *S. gigantea-invaded* plots (F_1,50_=3.29, *P*=0.0759; Table 2, Figure 1) but there were no significant effects of habitat type (Table 3, Figure 1).

### Two fungivorous nematode families benefitted from *S. gigantea-induced* increase in fungal biomass

The total nematode abundance, and density of the three fungivorous nematode taxa that were commonly present in both the Pleistocene sand and river clay locations were analysed to study the belowground impact of *S. gigantea* on the next trophic level of the soil food web. Representatives of the families Aphelenchidae, Aphelenchoididae and the genus *Diphtherophora* were used to determine whether and, if so, how the observed increase in biomass of higher fungi, and an unchanged fungal DNA levels are reflected in the local fungivorous nematode community.

Dominance by giant goldenrod did not affect the total nematode density (Table 2, Figure 1). Total nematode density (per 100 g dry soil) only differed significantly between habitats. The riparian sites had an estimated mean nematode abundance about two times higher than in semi-natural grassland soils (F_1,6_= 26.69, *P*= 0.0021; Table 2). Both Aphelenchidae (F_1,50_= 9.96, *P*= 0.0027) and Aphelenchoididae (F_1,50_= 8.44, *P*= 0.0054; Table 2, Figure 1) were more abundant in *S. gigantea-invaded* plots than in un-invaded plots. A significant interactive effect between habitat type and invasion status (F_1,50_= 6.92, *P*=0.0113) was observed for Aphelenchoididae indicating that the response to invasion was dependent on habitat type (Table 2) whereas this interactive effect between habitat type and invasion status was not significant for Aphelenchidae (F_1,50_= 3.16, *P*= 0.0814; Table 2).

For both fungivorous nematode taxa, the effect of *S. gigantea* was only seen in the riparian habitats. As compared to the un-invaded plots, Aphelenchidae densities were three times higher in invaded plots in riparian habitats (F_1,50_=11.30, *P*=0.0015; Table 3, Figure 1). Similarly, the estimated densities of Aphelenchoididae were around four times higher in *S. gigantea-invaded* riparian plots as compared to the non-invaded neighbouring plots (F_1,50_=14.23, *P*=0.0004; Table 3, Figure 1). Giant goldenrod stands did not affect the abundance of representatives of the genus *Diphtherophora* (Tables 2 and 3, Figure 1).

### Fungal indicator taxa associated with invasive *S. gigantea*

Invasion by *S. gigantea* resulted in a local increase in fungal biomass, but not in total fungal DNA. This remarkable observation was investigated in more detail by comparing the composition of the communities. As a crude measure for fungal DNA content we compared the number of primary reads per sample. Whereas soil samples from uninvaded plots gave rise to ≈ 95,000 (SD 39,000) reads per samples, on average ≈ 102,000 reads (SD 37,000) were generated from samples from invaded plots. No significant difference in number of reads per sample were found between invaded and non-invaded plots. Although it is hard to compare qPCR data with Illumina reads, the sequencing data confirm the absence of a difference in fungal DNA contents between *S. gigantea-invaded* and non-invaded plots.

PERMANOVA on Bray-Curtis dissimilarity profiles identified ‘habitat type’ (riparian vegetation *vs*. natural grassland) as the main factor responsible for the difference in fungal composition (Table 4). This factor explained ≈ 27% of the overall variance. The second most informative variable was ‘study site’ with an R^2^ value of 0.23. This is the variation in fungal communities between the various sampling sites within a habitat type. Against the substantial background variation caused by habitat type and study site, still a clear invasion effect could be discerned. Evidently, this effect explained a relatively low percentage of the overall variation, 1.7%, but this contribution was highly significant. As can be seen in Table 4, analyses of fungal communities for the two habitat types separately resulted in significant effects. It is noted that effect of plant invasion on fungal assemblages was more pronounced in the semi natural grasslands (*P*=0.001, and *P*=0.01 for riparian vegetation).

LEfSe (Linear discriminant analysis Effect Size) analysis allowed us to determine which fungal taxa contribute most to observed differences between *S. gigantea* invaded and un-invaded plots. With an LDA threshold of >2 the families Cladosporiaceae, Teratosphaeriaceae (both Ascomycota), Glomeraceae (Glomeromycota) and Kondoaceae (Basidiomycota) were shown to be more abundant in plots invaded with *S. gigantea* (Figure 2A). Analysis per habitat type revealed that only the Cladosporaceae were present in higher densities in both habitat types in invaded plots (Figure 2B and 2C). A higher abundance of members of the family Cucurbitariaceae (Ascomycota) was shown to be characteristic for the non-invaded plots.

**Figure 2.**
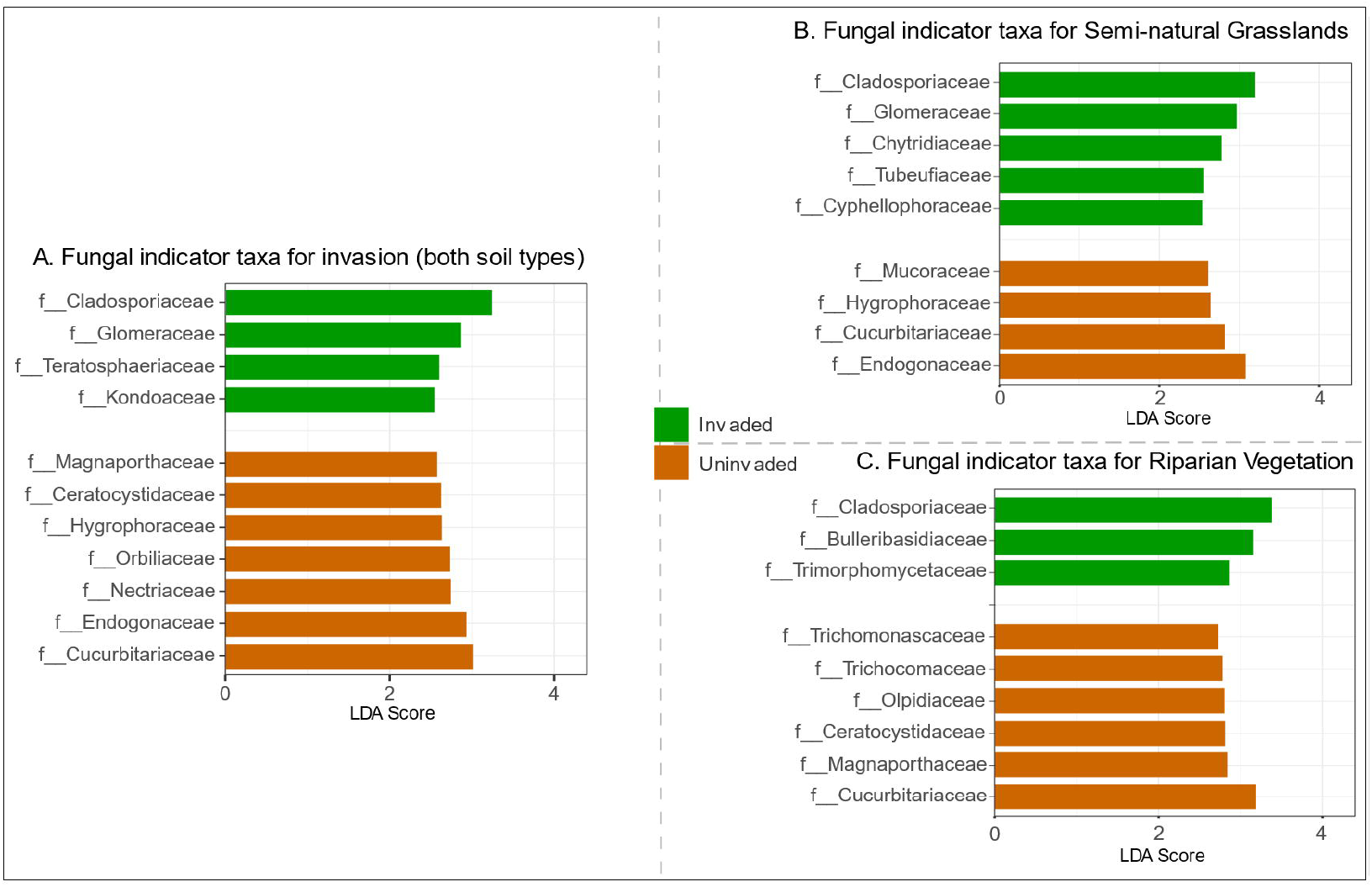
Discriminant fungal families indicated by LEfSe analysis (LDA threshold of 2) resulting from invaded (green) and uninvaded (brown) soils by Solidago gigantea.

**Table 4:** Summary PERMANOVAs on Bray-Curtis dissimilarity profiles of the fungal biome for the main effect and each habitat type. The effects of the following variables on the quantitative taxonomic composition of fungi were tested: Habitat type (Riparian Vegetation/Semi natural grassland), Study site (RV=3 SG=5), Invasion (Invaded/Uninvaded) and the interactions between Invasion and Habitat Type or Study Site. Differences are considered significant if P <0.01. P = probability associated with the Pseudo F statistic. Significant P values in bold.

|  | Main effect |  | Riparian Vegetation |  | Semi-natural grassland |  |
| --- | --- | --- | --- | --- | --- | --- |
|  | R2 | P | R2 | P | R2 | P |
| Habitat Type | 0.225 | **0.0001** |  |  |  |  |
| Study site | 0.237 | **0.0001** | 0.257 | **0.0001** | 0.354 | **0.0001** |
| Invasion | 0.018 | **0.0024** | 0.037 | **0.0050** | 0.032 | **0.0001** |
| Study site * Invasion | 0.033 | 0.3964 | 0.032 | 0.4252 | 0.053 | 0.2181 |
| Habitat * Invasion | 0.009 | 0.0523 |  |  |  |  |

## Discussion

The aim of this research was to characterize quantitative and qualitative shifts in the fungal community brought about by the invasive plant species *S. gigantea*. To assess the impact of this invasive plant species on below-ground fungal biomass, two kinds of components of the fungal cell membranes (either ergosterol or the fatty acid 18:2ω6) have been used. Consistently, *i.e*. during multiple growth stages, over multiple years, and at multiple location invasive giant goldenrod was shown to induce a local increase in fungal biomass (Quist et al. 2014; Stefanowicz et al. 2016). However, both a qPCR-based approach and an Illumina-based characterisation of fungal communities pointed at the absence of a quantitative shift in total fungal biomass. The observed *S. gigantea*-induced local increase in fungal biomass combined with the unaltered presence of fungal DNA prompted us to suggest that invasion of *S. gigantea* locally induced an increase in fungal biomass:DNA ratio. Qualitative characterization of the fungal assemblages revealed that *S. gigantea* invasion was accompanied by a local increase in abundance of members of the families Cladosporiaceae and Glomeraceae, and a decreased presence of Cucurbitariaceae.

To further investigate these apparently contradictory results, we focussed on representatives of the next trophic level; three fungivorous nematode taxa that were commonly present in both soil types. The densities of the fungivorous Aphelenchidae and Aphelenchoididae increased in *S. gigantea*-invaded plots, while the abundance of the members of the fungivorous genus *Diphtherophora* did not change. Moreover, we suggest that distinct food preferences explain why only for two out of three commonly present fungivorous nematode lineages an increase in density was observed. Arguments underlying this interpretation of our results are presented below.

### Apparent discrepancy between results from independent fungal biomass markers

Notably, the observed increase in ergosterol in *S. gigantea*-invaded plots was not accompanied by a comparable local augmentation of the total fungal DNA (Fig.1). In case of the phylum Ascomycota, a trend towards more rDNA in invaded plots was detected (Table 3, P= 0.0748). With regard to the Basidiomycota, it is noted that the apparent low density (Table 3) might be an underestimation as relatively high rDNA copy representatives were used to generate the calibration lines (Harkes et al. 2017; Lofgren et al. 2019).

Ergosterol is a frequently used marker for the assessment of fungal biomass in soil. It is sterol found in all Ascomycota and most Basidiomycota. Several representatives of the Zygomycota harbor ergosterol in their membranes as well, but this sterol is absent in the more basal fungal lineages (Weete et al. 2010). Using cultures of 6 non-basal fungal species (Montgomery et al. 2000) showed a tight correlation between ergosterol content and fungal biomass. Provided that local fungal community was dominated by later diverging divisions such as Ascomycota and Basidiomycota (as suggested by e.g. (Hannula et al. 2017)), ergosterol could be a reliable marker for fungal biomass.

With regard to the use of rDNA as a marker for fungal biomass, it should be mentioned that a genome-based survey revealed considerable variation estimation in rDNA copy numbers. Nevertheless, some phylum-specific characteristics have been observed. The average number of rDNA copies for Ascomycota is around 50 and shows limited variation. Basidiomycota harbor about twice as many rDNA copies, and this is accompanied by substantial variation among its members (Lofgren et al. 2019). Hence, rDNA copy numbers can only be used to assess fungal biomass in case there are no major differences between the community composition of the samples, and rDNA-based estimation of the Ascomycota is likely to be more accurate than the estimation of the Basidiomycota biomass.

Hence, although both ergosterol and rDNA copy number have their limitations as fungal biomass markers in soil, comparison of data from adjacent plots from the same habitat are probably valid. Fungi are known to be more flexible with regard to the biomass:DNA ratio than many other organismal groups (e.g. (Griffiths et al. 1997)). This is the result of the flexible cellular organization of fungi. The hyphal compartmentalization of fungi might be impaired by the partial or complete removal of septa, cross walls separating the fungal cells (Roper et al. 2011). Hence, growth of the mycelial network does not necessarily be accompanied by a comparable increase in the number of nuclei. Therefore, the difference in outcome between the two types of markers (biochemical or DNA-based) might be attributable to an increase in the fungal biomass:DNA ratio. Further research is required to investigate this hypothesis.

### The habitat (in)dependent impact of *S. gigantea* on fungivorous nematode lineages

Due to the apparently contradictory results obtained by the two types of fungal biomass markers, the effects of *S. gigantea* on major representatives of the next trophic level, fungivorous nematodes, were checked. Two out of three lineages of fungivores present both in the riparian zone and in the semi-natural grassland sites, Aphelenchoididae and Aphelenchidae, were shown to be stimulated in the presence of giant goldenrod, whereas the third lineages, the genus *Diphtherophoridae*, was unaffected by this invasive plant species. Moreover, the effect in the riparian habitats was much more pronounced than the effects in the sandy locations.

While the increase in Aphelenchoididae as a results of giant goldenrod invasion was found previously (Quist et al. 2014), the increase in Aphelenchidae was new. This difference may be explained by the seasonal fluctuations of nematode densities. Bulk soil concentrations of Aphelenchidae, Aphelenchoididae, and *Diphtherophora* were shown to have a distinct, taxon-dependent seasonality (Vervoort et al. 2012). In this research, samples were collected in late autumn, while samples analyzed in (Quist et al. 2014) were collected in early autumn. Hence it is conceivable that the *S. gigantea-induced* increase in Aphelenchoididae is only noticeable late in the season.

A pronounced boost of Aphelenchoididae and Aphelenchidae was observed in river clay soil, and a non-significant increase in sandy soils (e.g. Fig.1). This difference in response might be explainable by the soil texture-dependent species representation for each of these two families. Within the family Aphelenchoididae, the genus *Aphelenchoides* is dominant, and it comprises >100 predominantly fungivorous species. At least 30 species have been described for the constituting genera of the family Aphelenchidae, *Aphelenchus* and *Paraphelenchus*. No data on soil texture preference are available for *Aphelenchus, Paraphelenchus* or *Aphelenchoides*. We hypothesize that the species composition of families Aphelenchidae and Aphelenchoididae differed between the two main habitats. Apparently, the *Aphelenchoides* species present in the river clay soil could benefit more from the local increase in fungal biomass, than the *Aphelenchoides* species present in the sandy soils. We propose the same line of reasoning for the Aphelenchidae genera *Aphelenchus* and *Paraphelenchus*.

Hence, apparently contradictory results with regard to the impact of the invasive plant species *S. gigantea* on the fungal community, translated in a paradoxical effect on the fungivorous nematode community. In fact, only members of the families Aphelenchoididae and Aphelenchidae in riparian habitats benefitted from the presumed increase in fungal biomass. Representatives of the genus *Diphtherophoridae* were unaffected by the presence of giant goldenrod. We hypothesize that this difference in response could be caused by a difference in food preference between the lineages (Li et al. 2014; Okada and Kadota 2003; Ruess et al. 2000). It is noted that the fungivorous nematode densities reported in this study are relatively low. This could be a late season sampling effect. This effect has little impact on the current analyses as differences between uninvaded and invaded plots are considered rather than absolute changes.

### Effect of habitat-characteristic abiotic differences between habitat-type

Despite the differences in floristic composition, soil type and land use history between the riparian zone and the semi-natural grasslands (Quist et al. 2014), the overall biotic impact of giant goldenrod induced similar overall invasion effects. Nevertheless, Basidiomycota were more abundant in semi-natural grasslands than in the riparian zones (P= 0.017, Table 2). This could relate to a substantial pH difference. Whereas riparian zones had a relative neutral pH of 7.5, semi-natural grasslands had a nearly 2 units lower pH. As compared to bacteria, pH windows for optimal growth are wider for fungi (Rousk et al. 2010). In a more recent study, Zhang et al. (2016) investigated fungal communities in arctic soils with a pH range of over 2.5 units. In the most acidic sites, Basidiomycota showed a higher relative abundance as compared to sites with more basic soils. It is noted that the higher abundance in Basidiomycota in semi-natural grasslands did not result in a significant change in either of the fungivorous nematode lineages.

### Fungal indicator taxa related to invasion with *S. gigantea*

As shown in Fig. 2, the Ascomycete family Cladosporiaceae was one of the main families responsible for the *S. gigantea-induced* shift in fungal community composition. A closer look at the Cladosporiaceae OTUs revealed that *Cladosporium* was the dominant genus within this family.

*Cladosporium* is a fairly speciose genus, it comprises 189 species that are mostly saprotrophic but it also harbors some plant pathogens (Sandoval-Denis et al. 2016). Although in its native range leaves of showy goldenrod (*Solidago speciosa*) were shown to be infested by *Cladosporium asterum* causing brown rust pustules (website Missouri Botanical Garden (USA)), no information was found on *Cladosporium* being an important (Geisen et al.) pathogen of *S. gigantea* in Europe.

Recently (Koyama et al. 2019) studied root-inhabiting fungi in a wide range of native and exotic plant species in Canada. The plant selection included *Solidago canadensis*, in this context a native plant species. Ascomycota dominated the root-associated fungal communities, and within this division the Dothideomycetes – a class that includes the family Cladosporiaceae – were identified as the second most dominant class. From the overview of 27 plant species, the Cladosporiaceae abundances were in general negatively correlated with plant abundances, and the authors identified *Cladosporium delicatulum* as an endophytic plant pathogen to a substantial fraction of the plant species under investigation. In our research we focused on another *Solidago* species, outside its native range. In accordance with (Koyama et al. 2019) Cladosporiaceae was identified as an abundant fungal family in the two habitat types under investigation. However, an opposite, positive correlation between Cladosporiaceae and *Solidago gigantea* was observed in this study. Plant growth-promoting characteristics of a members of the genus *Cladosporium* could be marked as a possible benefit for *S. gigantea* associated with the fostering *Cladosporium* in its rhizosphere. Both *Cladosporium sphaerospermum* and *Cladosporium sp*. MH-6 were found to produced and release several types of gibberellins, which could explain their plant growth promoting characteristics (Hamayun et al. 2010; Paul and Park 2013).

Glomeraceae was the second most indicative family regarding the impact of *S. gigantea* on the local fungal community. Glomeraceae is a family of arbuscular mycorrhizal (AM) fungi and members of this family colonize the roots of a wide range of vascular plants including *S. gigantea* (Pirozynski and Dalpe 1989). (Vallino et al. 2006) characterized the AMF colonization of *S. gigantea* outside its native range, and identified *Glomus*, a genus belonging to the Glomeraceae, the dominant root colonizing AMF. This result underlines that invasive *S. gigantea* can recruit local AMF, and establish such a successful interaction that it ends up as one of the main fungal taxa typifying the community shift that was brought about by the invasive plant species.

In both habitat types, uninvaded plots were characterized by an increased presence of Cucurbitariaceae, just like the Cladosporiaceae a family that belongs to the class Dothideomycetes.

Little is known about the ecology of Cucurbitariaceae. Its members are known as saprobes on relatively recalcitrant organic materials such as wood, bark and leaves (Jaklitsch et al. 2018).

High-throughput sequencing revealed multiple fungal families as indicative for invasion. In the previous section we hypothesize that this difference in response could be caused by a difference in food preference between the lineages. Interestingly (Ruess et al. 2000) indicated *Cladosporium* as moderate feeding source for *Aphelenchoidides* sp, which is in line with our observations. Unfortunately, little is known about food preferences of different nematode taxa. Therefore, the indicator families observed in this research could be an interesting starting point for more targeted research to nematode feeding preferences. Especially the fungal families indicated for invasion in riparian habitat could be informative as Aphelenchidae and Aphelenchoididae showed a high significant increase in these soils.

## Conclusion

This study shows that *S. gigantea* invasion has a structural impact on the belowground soil community by increasing the fungal biomass independent of sampling moment, sampling year or habitat. The increase of fungal biomass is reflected in the next trophic level by a boost of two independent lineages of fungivorous nematodes, Aphelenchidae and Aphelenchoididae. Notably, this effect is more pronounced in the river clay soils in riparian zone than in the soil soils under the semi-natural grasslands. Nematodes show strong preferences for certain soil textures, even at species level. Therefore, we hypothesize that the different response levels might be contributable to differences of the species composition of Aphelenchidae and Aphelenchoididae fungivores in the two major habitats. Another fungivorous family, *Diphtherophoridae*, did not benefit from the local, *S. gigantea*-induced increase in fungal biomass.

The ergosterol-based observation of the increase in the fungal biomass a *S. gigantea*, could not be confirmed by DNA markers. Both qPCR-based assessment of total fungal DNA as well as the characterization of the fungal communities on the basis of variable 18S rDNA regions did not reveal a difference in fungal DNA contents between *S. gigantea*-invaded and non-invaded plots. The apparent discrepancy might be attributable to a change in DNA:biomass ratio. High throughput sequencing of the variable 18S rDNA regions V7-V8 revealed an increased abundance of Cladosporiaceae and Glomeraceae, and a decrease of the Cucurbitariaceae. Further investigation of the nature of these community shifts could further elucidate the change fungal DNA-biomass ratio as potentially provoked by the invasive plant species *S. gigantea*.

## Acknowledgment

We would like to acknowledge Natuurmonumenten, Staatsbosbeheer, Utrechts Landschap, the municipality of Wageningen and Reinaerde groenbeheer for allowing us to sample on their properties. We thank Herman Meurs (Unifarm, WUR) for his help during the OM content analysis and Eurofins Agro for analysing the soil C and N contents. This research was financially supported by the BE Basic Foundation, grant FS03.001.

## Author Contributions

M.T.W.V., C.W.Q. and J.H. were responsible for the experimental design. L.J.M.v.H., M.T.W.V. and C.W.Q. collected the soil samples. L.J.M.v.H. and S.J.J.v.d.E. isolated nematode DNA and performed the nematode qPCR assays. L.J.M.v.H. measured soil abiotic characters. M.H.M.H. and P.J.W.M. developed fungal qPCR primers. P.H. tested fungal qPCR assays. P.H., L.J.M.v.H. isolated bacterial and fungal DNA. P.H. performed the bacterial and fungal qPCR assays. P.H. performed the ergosterol measurements. G.G. performed the statistical analysis in SAS on the qPCR and ergosterol data. P.H. performed the two step PCR reactions in order to prepare the sequence library. J.J.M.v.S analysed the fungal sequence data and performed the statistical analysis. P.H. and J.H. wrote the manuscript; all others co-commented on the manuscript.

