## Supplemental Table 1 for "Characterization of the habitat- and season-independent increase in fungal biomass induced by the invasive giant goldenrod and its impact on the fungivorous nematode community"

**Supplementary Table 1:** Eight study sites located in riparian zones and in semi-natural grassland habitats are indicated below. Although ‘Blauwe Kamer’ is one riparian study site, samples were collected from two distinct areas within the nature reserve (1 and 2). Riparian zones are characterized by river clay soils, whereas the semi-natural grassland sites were located on Pleistocene sandy soils. Coordinates and years of *S. gigantea* introduction were obtained from Quist et al. (2014)

| **Habitat type** | **Study site** | **Soil type** | **Coordinates** | **Year of *S. gigantea* introduction** | **Number of plots pairs** |
| --- | --- | --- | --- | --- | --- |
| **Riparian zone** | Millingerwaard | River clay | 51° 51' 58.11'' N 6° 00' 35.47'' E | ~ 1950 | 8 |
|  | Ewijkse plaat | River clay | 51° 52' 47.36'' N 5° 44' 52.17'' E | ~ 1950 | 8 |
|  | Blauwe Kamer |  |  |  |  |
|  | *West* | River clay | 51° 56' 40.22'' N 5° 36' 19.90'' E | after 1950 | 4 |
|  | *East* | River clay and sand | 51° 56' 32.56'' N 5° 37' 09.54'' E | after 1950 | 4 |
| **Semi-natural grassland** | Dennenkamp | Pleistocene sand | 52° 01' 45.64'' N 5° 47' 53.50'' E | 1982 | 8 |
|  | Plantage Willem III | Pleistocene sand | 51° 58' 48.62'' N 5° 31' 08.47'' E | 1995 | 8 |
|  | Hollandseweg | Pleistocene sand | 51° 58' 49.89'' N 5° 40' 59.84'' E | before 2005 | 4 |
|  | Scheidingslaan | Pleistocene sand | 51° 58' 28.60'' N 5° 41' 55.40'' E | unknown | 4 |
|  | Reijerscamp | Pleistocene sand | 52° 00' 47.49'' N 5° 46' 08.64'' E | 2006 | 4 |
